# Syndecan-4 is required for VE-Cadherin trafficking during pathological angiogenesis

**DOI:** 10.1101/736553

**Authors:** Giulia De Rossi, Maria Vähätupa, Enrico Cristante, Sidath E. Liyanage, Ulrike May, Laura Pellinen, Saara Aittomäki, Zuzet Martinez Cordova, Marko Pesu, Hannele Uusitalo-Järvinen, James W. Bainbridge, Tero A.H. Järvinen, James R. Whiteford

## Abstract

New blood vessel formation, or angiogenesis, is characteristic of chronic diseases such as cancer, rheumatoid arthritis and vision-threatening conditions. Vascular Endothelial growth factor (VEGFA) and its receptor VEGFR2 drive neovascularization and hyperpermeability in these pathologies. One consequence of VEGFR2 activation is decreased stability of endothelial cell (EC) junctions through internalization of VE-Cadherin, allowing re-arrangement of sprouting ECs. Evidence suggests roles for heparan sulfate proteoglycans in angiogenesis and we show that Syndecan-4 (SDC4) expression is upregulated during pathological angiogenesis and is required for efficient VE-Cadherin internalization. Angiogenic responses in both tumor and neovascular eye disease models are impaired in Syndecan-4 null mice (Sdc4-/-), as is dermal hyper-permeability response to VEGFA. We show SDC4 resides at EC junctions and interacts with VE-Cadherin, an association lost upon VEGFA-stimulation, and this is SDC4 phosphorylation-dependent. Finally, we show that pathological angiogenic responses can be inhibited in a model of age-related macular degeneration by targeting SDC4. This study identifies SDC4 as a key component of VE-Cadherin trafficking and, as such, a critical regulator of pathological angiogenesis and vascular permeability.

## Introduction

Angiogenesis is a critical developmental process and is an essential component of physiological tissue regeneration (Eming et al, 2014). In cancer, inflammatory disorders and neovascular eye diseases, new blood vessel formation is a key feature in the pathogenesis and significant efforts have been made to control this process for therapeutic benefit (Adams & Alitalo, 2007; Amadio et al, 2016; Carmeliet & Jain, 2011; Jayson et al, 2016). The pro-angiogenic cytokine vascular endothelial growth factor A and its receptor VEGFR2 regulate angiogenesis in these pathologies and drugs targeting this axis have been used with substantial success in the clinic. However, there are issues of patient non-response, resistance to therapy and several side effects that are associated with VEGFA/VEGFR2 targeting drugs (Ebos et al, 2009; Kurihara et al, 2012; Meadows & Hurwitz, 2012), highlighting the need for better understanding of the underlying molecular basis of pathological angiogenesis.

VEGFA plays a key role in angiogenesis by regulating EC survival, proliferation and migration (Olsson et al, 2006). Junctional protein dynamics are key to EC re-arrangements during collective cell migration and thus their fine-tuned regulation is essential during angiogenesis (Dorland & Huveneers, 2017; Ingber, 2002). One of the major proteins residing at EC adherens junctions is VE-Cadherin, which forms homophilic interactions between neighboring ECs via its extracellular domain and provides a linkage to the actin cytoskeleton via α-, β- and p120 catenin, plaktoglobin and vinculin (Dejana, 2004). VEGFA controls EC junctional disassembly by promoting the β-arrestin-dependent endocytosis of VE-cadherin, resulting in disruption of endothelial barrier integrity and hyperpermeability (Gavard & Gutkind, 2006). Studies have shown that low VE-Cadherin expression or, alternatively, expression of a non-functional form of VE-Cadherin in ECs leads to incorrect cell rearrangements in newly-formed angiogenic sprouts (Bentley et al, 2014; Sauteur et al, 2014). Interestingly, pathological blood vessels, such as those observed in tumors and eye diseases, are characterized by abnormal morphology and functionality: tortuous, thin-walled, with diminished pericyte coverage and also hyperpermeable (Duh et al, 2017; Jain, 2005). There is considerable evidence to suggest a role for heparan sulfate proteoglycans in angiogenesis, in particular the syndecan family. The syndecans are a four-member family of single-pass transmembrane HSPGs with diverse roles in cell adhesion, receptor trafficking and growth factor signaling (Couchman, 2010). Recent studies have shown a central role for syndecan-2 (SDC2) in VEGFR2 signalling owing to enhanced 6-O-sulphation on its HS chains which promotes VEGFA binding (Corti et al, 2019). This work also highlighted that SDC4, the family member most closely related to SDC2, has no impact on VEGFA signalling and, in common with previous studies (Echtermeyer et al, 2001; Ishiguro et al, 2000b; Johns et al, 2016), that the *Sdc4-/-* mouse developed normally. It has been reported, however, that adult *Sdc4-/-* mice exhibit defects in the development of the foetal labyrinth and in granulation tissue formation after wounding (Echtermeyer et al, 2001; Ishiguro et al, 2000a) which is suggestive of a functional requirement for SDC4 in new blood vessel formation.

Here, we set out to elucidate the function of SDC4 in angiogenesis using a variety of disease models. We found that *Sdc4-/-* animals are protected in both murine tumor models and models of neovascular eye disease, that SDC4 expression is upregulated during pathological angiogenesis in the mouse but not during development and is selectively expressed on immature blood vessels in retinas of diabetic retinopathy patients. We show that SDC4 is required for efficient VE-Cadherin trafficking away from EC junctions and demonstrate that phosphorylation of the SDC4 cytoplasmic domain is a critical step in this process. Finally, we show that targeting SDC4 in an *in vivo* model of neovascular AMD ameliorates the outcome. These findings identify SDC4 as an important regulator of VE-Cadherin trafficking in VEGFA-driven junctional reorganization during pathological angiogenesis, and as such has the potential to be targeted for therapeutic benefit.

## Results

### Tumor angiogenesis is impaired in Sdc4-/- animals

The formation of new blood vessels is an essential component of tumor growth and development (Jain, 2014). To investigate whether SDC4 plays a role in tumor angiogenesis, we employed a skin epidermal carcinogenesis model (2-stage DMBA/TPA), the progression of which is angiogenesis-dependent (Filler et al, 2007; Perez-Losada & Balmain, 2003), to measure tumor angiogenesis in WT and *Sdc4-/-* mice. At the end point of 19-weeks, WT animals had on average 2.4 times more tumors than *Sdc4-/-* mice and the incidence of large tumors was 6-fold higher (**Fig. 1A-D**). In the DMBA/TPA model, tumor angiogenesis takes place first by increased blood vessel density and later, during papilloma formation, by increased blood vessel diameter (VEGFA-driven process). Histochemical staining for the EC marker CD31 revealed smaller blood vessel lumens in tumors formed in *Sdc4-/-* mice compared to WT (**Fig. 1 E and F**), suggesting that lack of SDC4 retards tumor angiogenesis. To dissect whether this phenotype is due to *Sdc4-/-* tumor cells or *Sdc4-/-* tumor vasculature, we injected B16F1 melanoma cells, which express SDC4 (**Fig. S1A**), into the flank of WT and *Sdc4-/-* mice. Mice were sacrificed after 2 weeks and both tumor volume and weight were greatly reduced in *Sdc4-/-* animals when compared to WT controls (**Fig. 1G and H, Fig. S1B**). Critically, histological analysis of tumor sections from WT animals revealed vascular structures with, in many cases, a well-defined lumen (white arrows in **Fig. 1I**) while in *Sdc4-/-* mice-derived tumors, ECs failed to organize into tubules and were more sparsely distributed (**Fig. 1I and J**).

**Figure 1:**
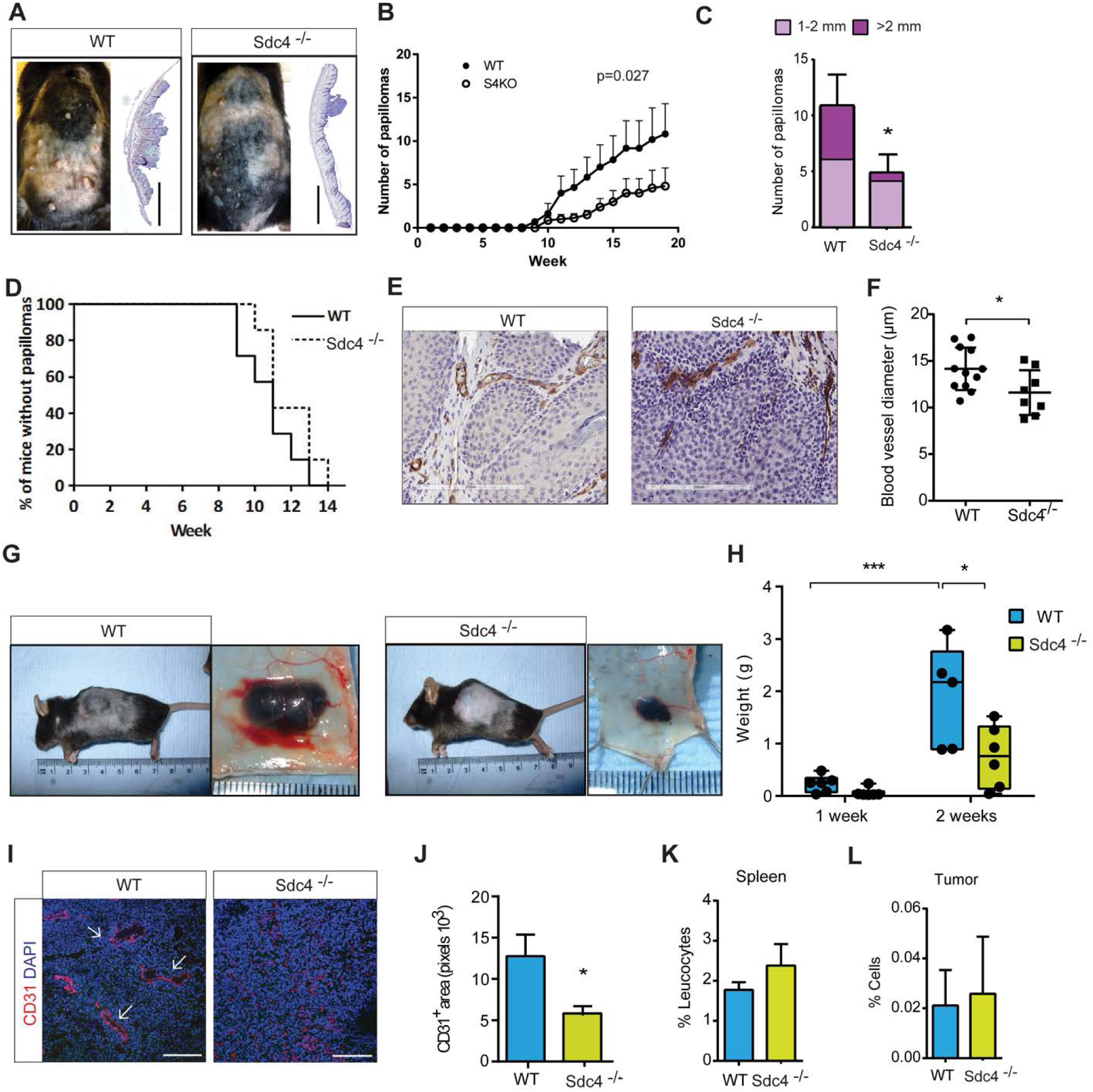
Tumor angiogenesis is impaired in *Sdc4-/-* mice. **(A)** Papilloma formation is reduced in *Sdc4-/-* mice in the DBMA/TPA model. Micrographs of animals (left) and sections of skin (right, H&E stained) from WT and *Sdc4-/-* animals at week 19 (scale bar = 2 mm). **(B)** Number of large tumors (≥2 mm/mouse) over time and **(C)** size of papillomas at end of the experiment (n=7 mice/group). **(D)** Time-course of papilloma incidence in WT and *Sdc4-/-* mice. The percentage of tumor-free animals at each time point is shown. Papillomas were observed in WT and *Sdc4-/-* mice at 9 weeks and after week 12 none of the animals was tumor-free. Because the proportional hazards assumption appeared correct, a survival plot was generated and analyzed via log-rank (Mantel-Cox) test. **(E)** Tumor sections from WT and *Sdc4-/-* animals were immuno-stained for the EC marker CD31and vessel width was measured. **(H)** Vessels from *Sdc4-/-* papillomas were narrower. **(G)** Micrographs of B16F1 melanomas from WT and *Sdc4-/-* animals showing reduced tumor volume as quantified in **(H)** (n=5-6 mice/group). **(I)** Tumor vessels (arrowheads) appear in WT sections but are not obvious in B16F1 melanomas from *Sdc4-/-* mice (Ki-67, blue; CD31 red, scale bar = 100 µM), **(J)** quantification of tumor vessel coverage (n=5/group, 3 images/animal). *P < 0.05. Error bars indicate SEM. Levels of NK cells are equal in WT and *Sdc4-/-* animals in both spleen **(K)** and B16F1 tumor immune infiltrates **(L)** (n=3 mice/group).

Reduced tumor volumes in the Lewis lung carcinoma model in *Sdc4-/-* animals has been reported and this correlated with an increased number of Natural killer (NK) cells in immune infiltrates from tumors (El Ghazal et al, 2016). This prompted us to analyze the immune infiltrates in our tumor models by flow cytometry. We found no differences in B16F1 tumors in numbers of NK cells between WT and *Sdc4-/-* animals (**Fig. 1K and L**) and this was also true of other leukocyte subsets including B cells, T cells, monocytes and neutrophils (**Fig. S2A and B**). Analysis of skin and tumors from the DMBA/TPA model also showed no trend for differential inflammatory responses, i.e. macrophages, T cells or neutrophils, between WT and *Sdc4-/-* animals and this was confirmed both by immuno-histochemical analysis (**Fig. S3A-F**) and flow cytometry (**Fig. S4A-C**). We also quantified cell proliferation rate with the proliferation marker Ki67 in the epidermis of DMBA/TPA treated mice. No statistically significant differences between *Sdc4-/-* and WT animals could be detected either in epidermis or in papillomas (**Fig S5 A and B**) and this was also true of skin thickness (**Fig S5C and D**). These observations led us to believe that the reduced tumor growth in *Sdc4-/-* mice observed in both models was primarily due to defective tumor vascularisation.

### Angiogenesis in ocular disease models requires Sdc4

Having established that tumor angiogenesis was impaired in *Sdc4-/-* mice, we next tested whether other pathological angiogenic responses were affected in these animals. Oxygen-induced retinopathy (OIR) is a hypoxia-driven angiogenesis model that recapitulates features of diabetic retinopathy and retinopathy of prematurity, whereas, laser-induced choroidal neovascularization (CNV) models neovascular (‘wet’) age-related macular degeneration (Grossniklaus et al, 2010). OIR was performed on 7-day old WT and *Sdc4*-/- pups by exposing them to hyperoxia (75 % O2 for 5 days) leading to the abolition of the retinal vasculature, before being returned to normoxia which stimulates a neovascular response. Exposure to oxygen led to a comparable loss in retinal vasculature in both *Sdc4-/-* and WT control littermates (**fig. S6, A and B**); however, upon return to normoxic conditions it was evident that the formation of neovascular tufts (a hallmark of pathological angiogenesis) was significantly reduced (∼ 40% decrease) in the retinas of *Sdc4-/-* mice compared to WT mice (**Fig. 2, A and B**). In contrast, the physiological neovascular response associated with hypoxia in *Sdc4-/-* animals was only slightly increased as determined by smaller avascular areas in these animals 5 days post-hyperoxia compared to WT littermates (**fig. S6, C and D**). We next investigated whether neovascularization is also attenuated in *Sdc4-/-* mice in the laser-induced choroidal neovascularization (CNV) model. Laser photocoagulation was performed on the eyes of 6-week old WT and *Sdc4-/-* littermates to stimulate a neovascular response in the choroid. Areas of CNV measured 7 days after injury by fluorescein angiography were found to be significantly smaller in the eyes of *Sdc4-/-* mice compared to WT mice (**Fig. 2, C and D**). CNV lesions were predominantly composed of ECs on the basis of lectin GS-II staining and we confirmed the reduction in lesion volume (∼80% decrease) in *Sdc4-/-* mice using 3D confocal densitometry (**Fig. 2, E and F**). This data suggests that angiogenic defects in *Sdc4-/-* animals are not restricted to cancer models, but there is also a defect in the angiogenic response in models of disease in the eye.

**Figure 2:**
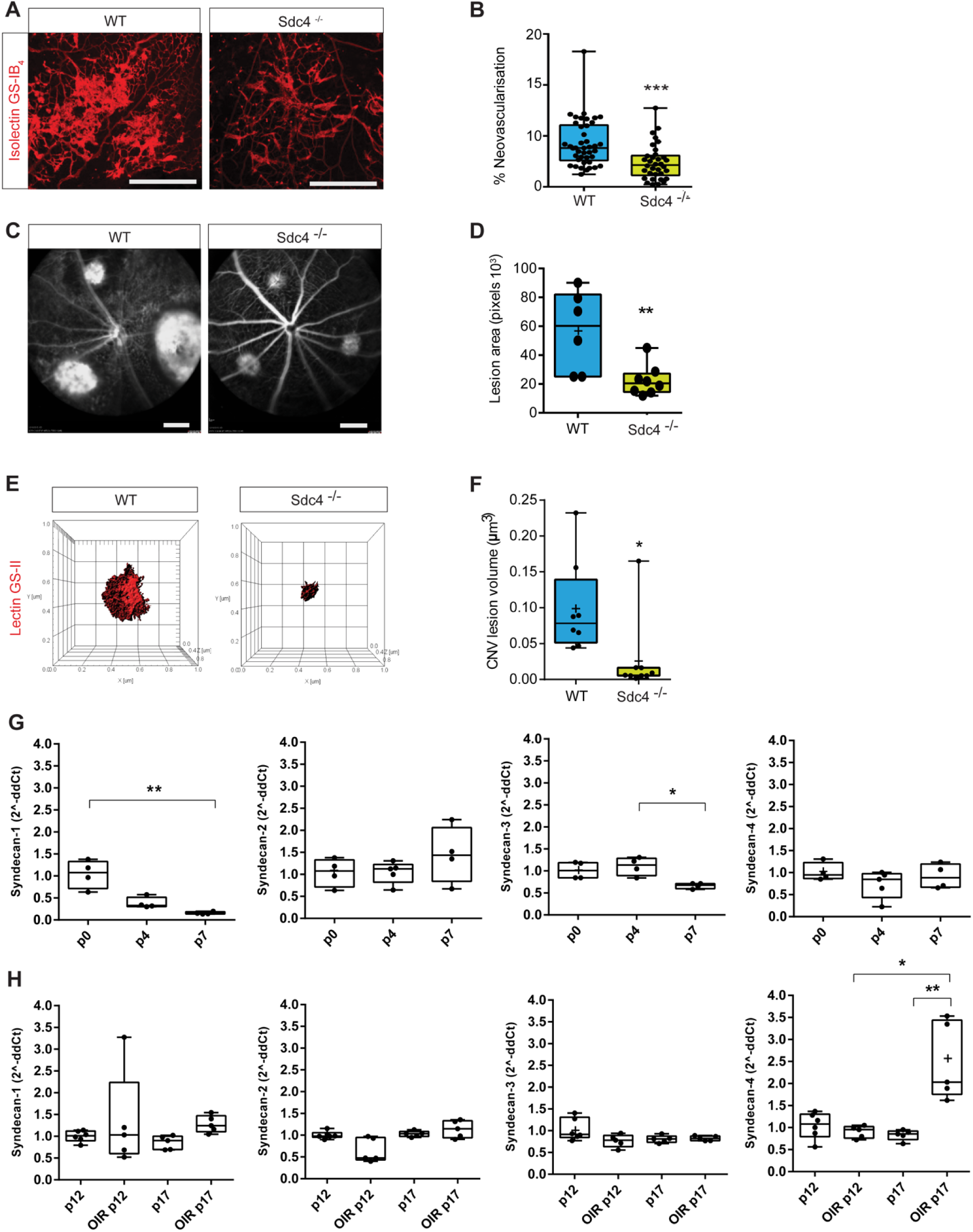
Pathological neovascularization in models of ocular disease is impaired in *Sdc4-/-* animals. **(A)** Micrographs showing the pre-retinal neovascularization response (stained with BS1-isolectin) to OIR in P17 WT neonates is greater than in equivalent *Sdc4-/-* animals (Scale bar = 500 µM). **(B)** Quantification of pre-retinal neovascularization (∼40 eyes/group). **(C)** Micrographs showing *Sdc4-/-* animals exhibit less angiogenesis in the laser induced CNV model as evident from reduced lesion area **(D)** (n=6-8 animals/group, Scale bar = 2.4 mm). **(E)** Staining of CNV lesions with lectin GS-II followed by 3D confocal reconstruction. **(F)** CNV lesion volume was calculated using Imaris Bitplane software on the basis of lectin GS-II staining. n=5-6 animals for each group. Centre line, median. Plus sign, mean. Error bars indicate min and max values. *P < 0.05; **P < 0.01; ***P<0.001. **(G)** Syndecan gene expression profile during early stages of murine retinal angiogenesis. Eyes were enucleated at postnatal day 0, 4 and 7, mRNA extracted and quantified by qPCR. **(H)** Syndecan gene expression in neonates subjected to OIR (P12 vaso-obliteration phase, P17 angiogenic phase) and in untreated controls. Centre line, median. Plus sign, mean. N=4-6 animals for each group. Error bars indicate min and max values. *P < 0.05; **P < 0.01.

### SDC4 expression is upregulated during pathological angiogenesis

We next sort to compare syndecan gene expression in both the context of developmental and pathologic angiogenesis in the murine retina. During the early stages of murine postnatal development, angiogenesis occurs in the eye leading to the formation of a superficial retinal vascular plexus and in C57BL/6 mice this happens during the first 8 days after birth. Quantitative rtPCR was performed on retinal mRNA from day P0, 4, and 7 neonates to measure the expression of *Sdc-1, -2, -3* and *-4* during development. *Sdc-1* and *-3* showed downregulated expression over time, whereas *Sdc-2* and *-4* remained relatively constant (**Fig. 2G**). Next, we examined the transcript profile of syndecans in a murine model of retinal pathological neovascularization, OIR. When pups were subjected to OIR, we detected a marked increase in *Sdc4* gene expression in retinas at P17, the time point at which neovascularization peaks, whereas the expression of the other 3 syndecans remained stable (**Fig. 2H**). Taken together, these data show that all four syndecans are expressed in the murine eye, but only SDC4 is upregulated during pathological angiogenesis in OIR retinas.

To investigate whether this was also the case in a human disease setting, we analyzed the expression of SDC4 in the retinal neovascular membranes that develop in human diabetic retinopathy patients. Neovascular membranes were collected from type I diabetes patients suffering from retinopathy, who had either already developed or had threatening tractional retinal detachment due to fibro-vascular proliferation. These tissue samples represented the end stage of the disease, but still contained regions with active pathological angiogenesis. Histological analysis of these samples revealed SDC4 expression predominantly associated with blood vessels (CD31^+^ areas). Moreover, immunostaining of adjacent human tissue sections for SDC4, VEGFR2 (a marker of pathological immature blood vessels) and CD31 revealed that SDC4 expression strongly correlates with CD31^+^/VEGFR2^high^ blood vessels, whereas only moderate expression was detected in CD31^+^/VEGFR2^low^ areas (**Fig. 3A to C**). We conclude from this that, at least in the context of neovascular eye disease, SDC4 expression positively correlates with new blood vessel formation during pathological angiogenesis in both murine models and in human disease.

**Figure 3:**
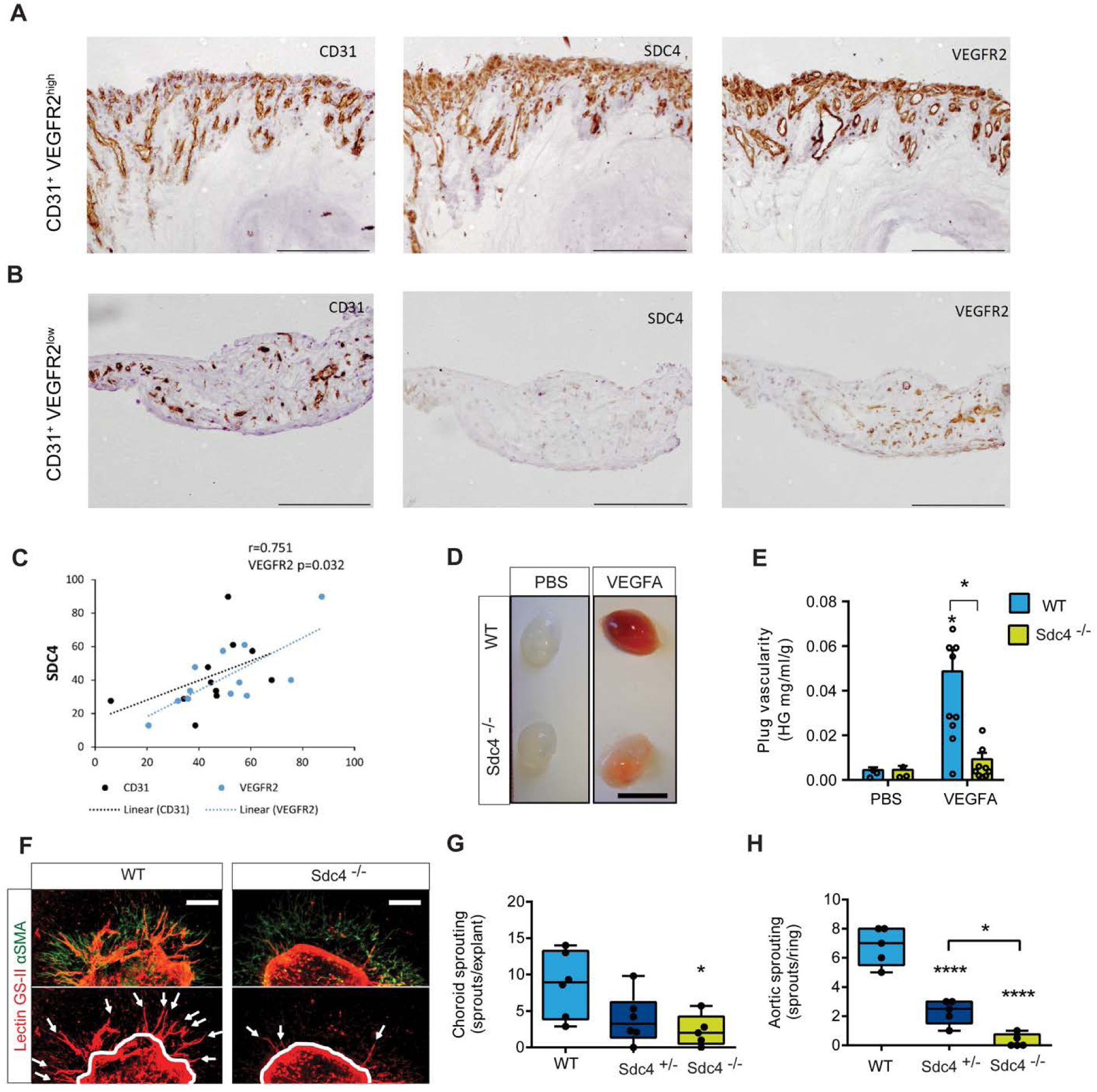
VEGFA stimulated angiogenic responses require *Sdc4*. **(A and B)** Consecutive sections of human proliferative diabetic retinopathy membrane stained for CD31 marker of blood vessels), VEGFR2 (marker of new immature vessels), and SDC4. Scale bar, 200 μm. Images are representative of n=6 patients. Correlation between SDC4, CD31 and VEGFR2 expression in neovascular membranes from diabetic patients **(C)**. Multiple linear regression analysis of correlation between SDC4 and CD31^+^ or VEGFR2^+^ area (%) (Correlation R=0.751, p=0.036, coefficients CD31=0.963, VEGFR2=0.032). n=6 patients (2-3 sections per patient), individual ROIs (n=11) were used for the analysis. **(D)** Matrigel supplemented with PBS or VEGFA (50 ng) was injected subcutaneously in the flank of WT or *Sdc4-/-* mice. Images are representative of plugs extracted 7 days post-injection. Scale bar, 1 cm. **(E)** Plug vascularity was expressed as the amount of hemoglobin released from the plugs per ml of Matrigel and normalized for the plug weight. n=5-6 animal for each group (2 plugs per animal). **(F)** Fragments (1 mm2) of choroid were dissected from adult WT, Sdc4 +/- and Sdc4-/- littermates and cultured as explants in VEGFA-containing medium. Explants were stained with lectin GS-II (red) and anti-αSMA (green) after 7 days in culture. Arrows show angiogenic sprouts. Scale bar, 10 µM. **(G)** Sprouting was quantified by manually counting the number of sprouts per explant. n=5-6 animal for each group, 10 explants per animal. **(H)** Quantification of sprout formation from aortic ring explants (n=5-6 animal for each group, 15-20 rings per animal).

Despite the profound phenotype that *Sdc4-/-* mice develop in pathological angiogenesis models, we did not observe any major difference in postnatal retinal vascular development between WT and *Sdc4-/-* mice at P6 (**Fig. S7, A and B**). However, a slight but significant reduction in vascularity was observed suggesting some involvement of SDC4 in developmental angiogenesis (**Fig. S7, C and D**) and this was also reflected in measurements of vascular area coverage **(Fig. S7E)** in which both *Sdc4+/-* and *Sdc4-/-* showed a reduction. Despite this, there were no differences in the number of arteries and veins (**Fig. S7F**). Analysis of additional adult vascular beds (skin, muscle and connective tissue) revealed that *Sdc4-/-* animals developed a normal vascular network (**Fig. S8, A to E**). Thus, SDC4 has a critical role in pathological neovascularization in the models tested, whereas its effects on developmental angiogenesis are more subtle.

### SDC4 is required for VEGFA-driven angiogenic responses

The principal driver of angiogenesis in the models described above is VEGFA, we therefore wanted to explore whether responsiveness to VEGFA was a factor in the phenotypes we observed in *Sdc4-/-* animals. WT and *Sdc4-/-* mice were injected with Matrigel containing VEGFA which triggered an angiogenic response in WT mice as evidenced by plug hemoglobin content. In contrast, an angiogenic response was notably reduced from VEGFA-containing plugs from *Sdc4-/-* mice (**Fig. 3, D and E**). We next adopted a more reductive approach in which tissue explants from both choroid membranes and aortas from *Sdc4-/-, Sdc4+/-* and WT littermates were embedded in a collagen I matrix and exposed to VEGFA. In both models, significantly more angiogenic sprouts emerged from WT, compared to *Sdc4-/-* explants (**Fig. 3, F to H**). Explants from heterozygous (*Sdc4+/-*) mice exhibited an intermediate phenotype indicating that the angiogenic response to VEGFA associated with SDC4 is subject to gene dosage effects. These data suggest that the pro-angiogenic role of SDC4 in pathological neovascularization is linked to VEGFA stimulation.

### Sdc4 has a role in VEGFA induced VE-Cadherin trafficking from EC junctions

We hypothesized that endothelial SDC4 could regulate VEGFA-dependent EC migration. We analyzed EC migration by light microscopy 6 hours after a longitudinal scratch was made on a confluent monolayer of cells. Results showed that while primary *Sdc4-/-* lung ECs were able to migrate in a growth factor-rich environment (FBS), albeit slightly less than WT ECs, their migratory response to VEGFA stimulation was negligible (**Fig. 4, A and B**).

**Figure 4:**
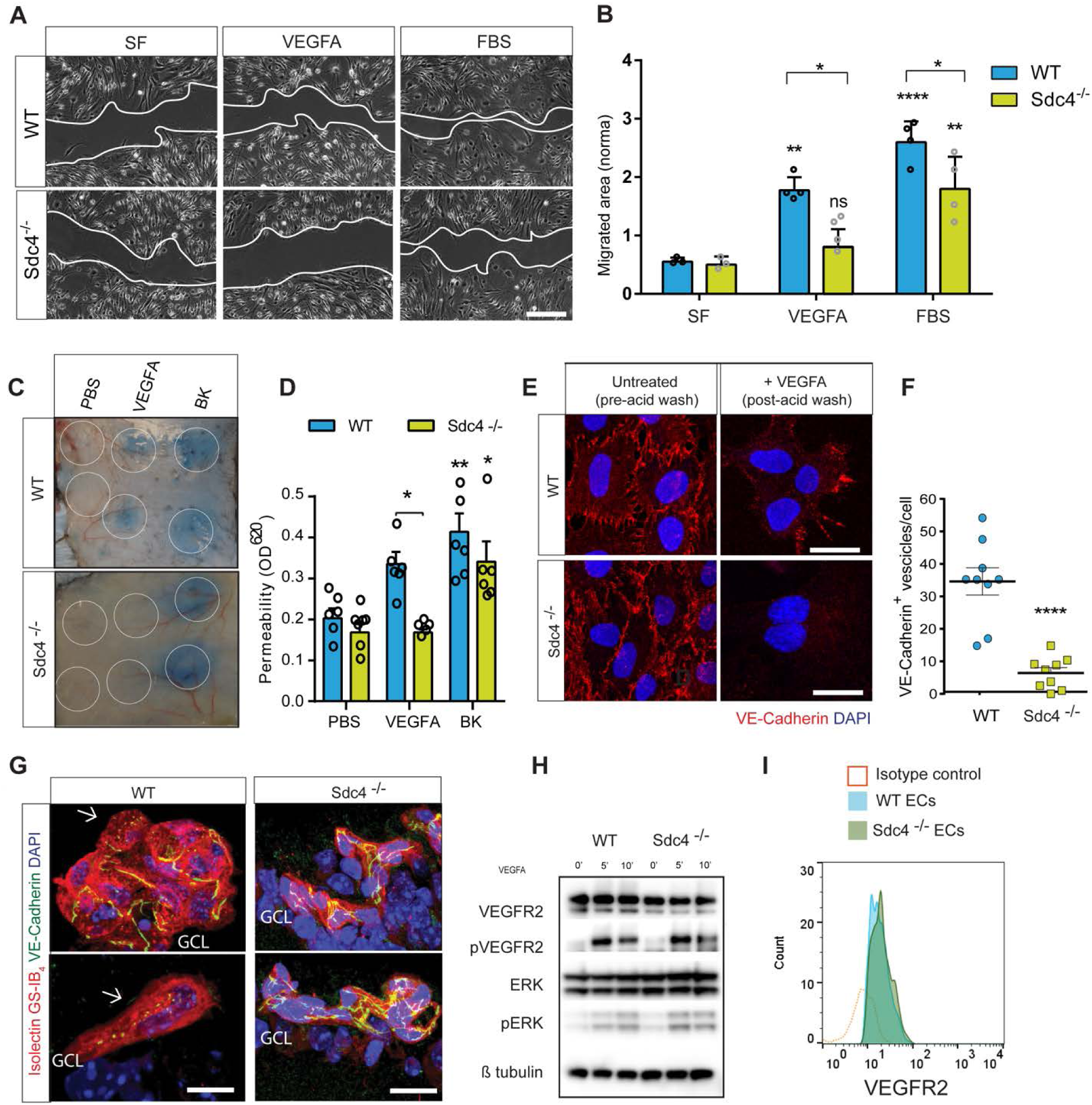
VEGFA induced EC junctional rearrangements require SDC4. **(A)** MLECs were scratched and incubated in serum-free (SF) medium or with VEGFA (100 ng/ml) or FBS (10%) containing medium for 16 hours. Phase contrast images showing final scratch area (white lines). Scale bar, 200 µm. **(B)** Migrated area was calculated by subtracting the final scratch area to the initial scratch area using ImageJ. n=3. **(C)** Evans blue was injected into the tail vein of WT and Sdc4-/- mice, followed by subcutaneous injections of PBS, VEGFA (100 ng) or Bradykinin (BK, 100 μg). Images show the local extravasation of the dye from underneath the skin 90 minutes post-injection, white circles indicate the approximate injection area. **(D)** Skin punches corresponding to the injection sites were collected and Evans blue extracted in formamide overnight. Permeability was quantified by measuring the optical density at 620 nm of the skin extracts and values were normalized on the basis of tissue weight. n=6-8 animals per condition. Data are mean ± s.e.m. *P < 0.05; **P < 0.01; ****P<0.0001. Statistical comparisons were made between PBS and treatments within the same genotype unless otherwise indicated. **(E)** MLECs from WT and *Sdc4-/-* mice were washed with PBS and incubated with anti-VE-Cadherin antibody at 4 °C for 1 hour. Unbound antibody was washed away and cells stimulated with VEGFA (30 ng/ml) for 10 min at 37 °C to promote VE-Cadherin internalization. Cells were then subjected to an acid wash to remove cell surface antibody. VE-Cadherin (red), DAPI (blue). Scale bar, 20 μm. **(F)** Images of MLECs treated with VEGFA and acid-washed were analyzed on Imaris software and VE-cadherin+ vesicles were counted and divided by the number of nuclei in the field of view. n=3. Images are representative of one experiment where 9 images per condition were analyzed. **(G)** Two representative images of WT and *Sdc4-/-* OIR retinas paraffin-embedded, sectioned and stained with isolectin GS-IB4 (red), VE-Cadherin (green) and DAPI (blue). Arrows indicate VE-Cadherin+ vesicles. GCL: ganglion cell layer. Blood vessels are pre-retinal tufts. Scale bar, 20 μm. VEGFR2 signaling is unaffected in Sdc4-/- primary MLECS. **(H)** Western blots of either WT or *Sdc4-/-* MLEC lysates harvested at different time points of showing a VEGFA stimulation. Levels of phoshpo-VEGR2, Erk, Src and VE-Cadherin were assayed. **(I)** Cell surface expression of VEGR2 is the same on WT and *Sdc4-/-* MLECs as measured by flow cytometry.

Angiogenesis and EC migration involve many changes to the endothelium not least the dissociation of EC junctions which leads to increased microvascular permeability *in vivo*. One way of modelling this process is to measure VEGFA-induced hyperpermeability using the Miles assay. Leakage of Evans blue from the dermal microvasculature was measured in response to VEGFA and bradykinin (a known vasodilator) in WT and *Sdc4-/-* animals. Vascular leakage in response to VEGFA was significantly reduced in *Sdc4-/-* animals (**Fig. 4, C and D**). A slight but insignificant reduction in vascular permeability was also observed in response to bradykinin. This led us to postulate that SDC4 could play a role in the regulation of VEGFA-induced VE-Cadherin trafficking and tested this hypothesis in an antibody-feeding assay. We found that ECs isolated from *Sdc4-/-* mice displayed reduced VE-cadherin internalization following 10 min exposure to VEGFA compared to WT ECs (**Fig. 4, E and F**). In line with this observation, newly-formed blood vessels in retinas retrieved from *Sdc4-/-* OIR mice showed modest presence of intracellular VE-cadherin^+^ vesicles compared to WT vasculature where VE-cadherin immunostaining is more discontinuous and VE-Cadherin^+^ vesicles are abundant (**Fig. 4G**).

Taken together these data could potentially be explained by inefficient or impaired VEGFA signaling via its interaction with VEGFR2. We ruled out this hypothesis by looking at changes in the phosphorylation status of VEGFR2 in WT and *Sdc4-/-* primary lung ECs and the key downstream kinases Erk1/2 where no differences were observed in response to VEGFA stimulation (**Fig. 4H**). We also compared cell surface expression levels of VEGFR2 by flow cytometry on these cells and found no difference (**Fig. 4I**). These findings confirmed that SDC4 does not act as a VEGFR2 co-receptor and were consistent with other studies suggesting that it is indeed SDC2 which mediates VEGFR2 signaling by binding VEGFA through its HS chains (Corti et al, 2019). This led us to examine an alternative hypothesis where the role of SDC4 is in fact not as a co-receptor but indeed downstream of VEGFA signaling.

### VE-Cadherin trafficking in response to VEGFA requires SDC4

We first performed immunofluorescence confocal microscopy on confluent HUVECs to look for endogenous SDC4 expression. A pool of SDC4 was evident at EC junctions colocalising with both VE-Cadherin and VEGFR2 (**Fig. 5A**). To better study the localization and trafficking of SDC4 in HUVECs, we generated an eGFP tagged form of SDC4 (eGFP inserted between I^32^ and D^33^ of murine SDC4 cDNA, **Fig. 5B**), transfected HUVECs and looked at the cellular localisation of Sdc4-eGFP. Consistent with **Fig. 5A**, a pool of eGFP-SDC4 localised to EC junctions (**Fig 5B**). Additionally, we could demonstrate that eGFP-SDC4 was expressed on the cell surface of HUVECs both by flow cytometry (**Fig. S9A**) and cell surface biotinylation assays and that western blotting of both total cell lysates and cell surface fractions for eGFP showed that this mutant was glycanated based on the presence of high molecular weight eGFP immuno-reactive smears (**Fig. S9B**). We could also show that overexpression of Sdc4-eGFP in HUVECS did not impact either cell surface expression of VEGFR2 or VEGFA-induced VEGFR2 phosphorylation or (**Fig. 5C and D**). Taken together these data suggest that expression of eGFP-SDC4 in HUVECs presents a viable tool for monitoring the cellular localisation of SDC4 in ECs.

**Figure 5:**
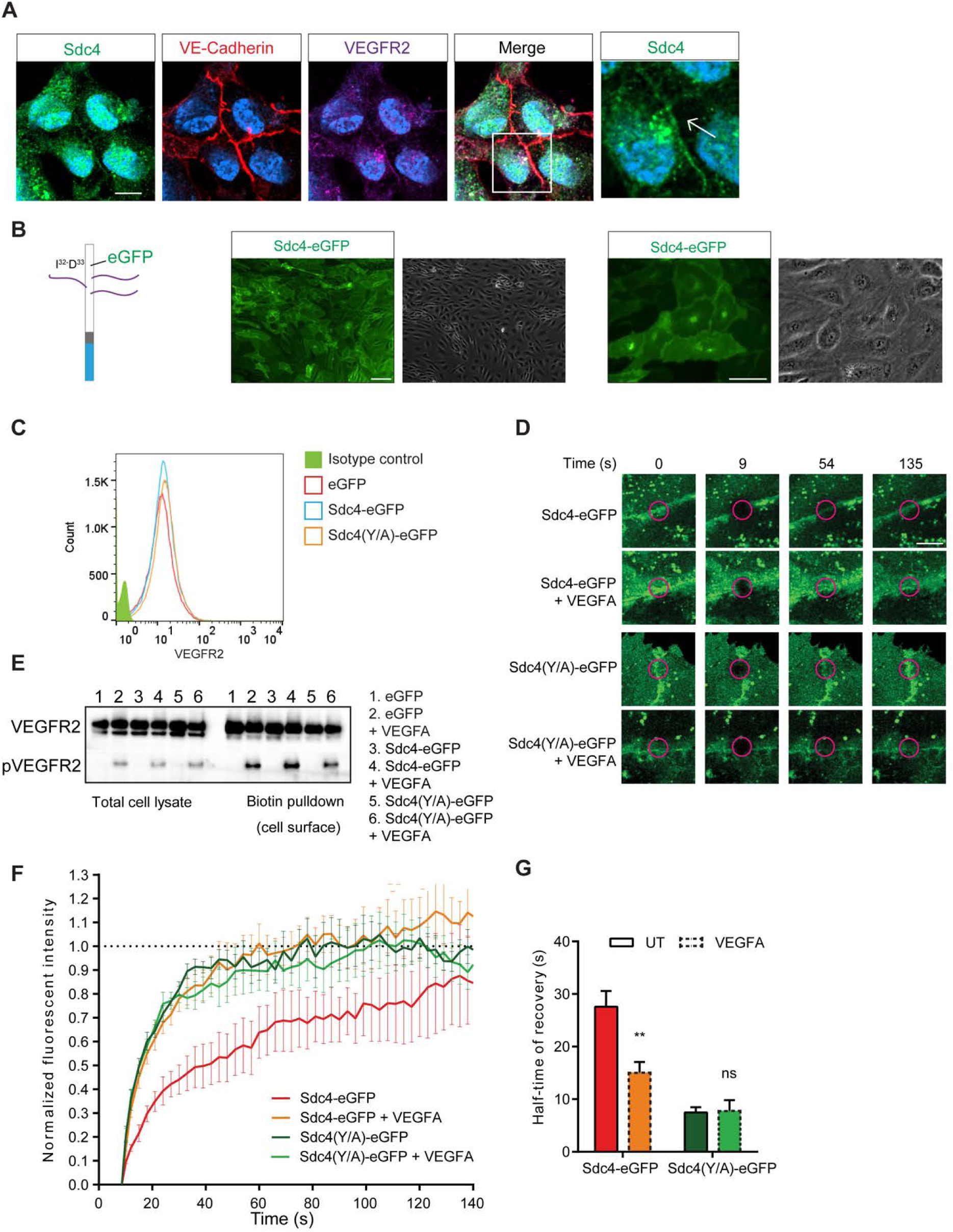
A pool of SDC4 resides at EC junctions and appear to be in a complex. **(A)** Immunofluorescence staining for endogenous SDC4 (green) shows co-localisation with VE-Cadherin (red) and VEGFR2 (purple) at EC junctions. Scale bar, 10 µm. **(B)** The complete coding sequence of eGFP was inserted into murine SDC4 between I^32^ and D^33^ and cloned into lentiviral expression vectors (diagram). Phase contrast and fluorescent images at x20 and x40 of transfected HUVECS are shown on the right and these confirm Sdc4-eGFP localizes to EC junctions. Scale bar, 100 µm. **(C)** Cell surface VEGFR2 expression in HUVECs is unaffected by overexpression of Sdc4-eGFP and Sdc4(Y/A)-eGFP as measured by flow cytometry. **(D)** Western blots showing that VEGR2 phosphorylation (Y^1175^) in response to VEGFA is not affected in HUVECs expressing Sdc4-eGFP and Sdc4(Y/A)-eGFP. **(E)** Confocal images of FRAP of HUVECs expressing Sdc4-eGFP and Sdc4(Y/A)-eGFP at the time points indicated in the presence or absence of VEGFA stimulation. **(F)** Quantification of fluorescence recovery of Sdc4-eGFP and Sdc4(Y/A)-eGFP and **(G)** plots of the half-life of recovery for each of the treatments.

Fluorescence recovery after photo-bleaching (FRAP) is a technique used to measure the kinetics of diffusion of a particular fluorescent molecule in living cells. We performed photobleaching of Sdc4-eGFP at EC-EC contacts and observed the rate of recovery of fluorescence both under basal and VEGFA-stimulated conditions. Recovery of eGFP-SDC4 fluorescence was significantly enhanced by the addition of VEGFA supporting the idea that SDC4 is acting downstream of VEGFA in some capacity (**Fig. 5E-F**). Previous studies have shown that phosphorylation of SDC4 at Y^180^ by Src kinase is essential for trafficking of integrins in fibroblasts (Morgan et al, 2013). To this end, we asked whether phosphorylation of this residue was important for the VEGFA stimulated response observed above with Sdc4-eGFP. We generated a mutant form of Sdc4-eGFP in which Y^180^ of SDC4 was mutated to an Alanine (Sdc4(Y/A)-eGFP). We transfected this into HUVECs and verified it localises to EC-EC contacts, is expressed on the cell surface, does not impact on VEGFR2 expression or signaling and was glycanated (**Fig. 5C and D, Fig S9 A and B**). Interestingly, fluorescence recovery of this non-phosphorylatable form of eGFP-SDC4 was substantially more rapid under basal conditions and VEGFA stimulation had no effect on this (**Fig. 5D-F**). One way of interpreting this data is that VEGFA stimulation results in the disruption of a complex involving SDC4 meaning that it can more rapidly diffuse back into photobleached areas. Participation within this complex is dependent on SDC4 phosphorylation since the non-phosphorylatable Y^180^ mutant recovers more rapidly.

Given the defect in VE-Cadherin internalization we observed in the *Sdc4-/-* animals and ECs, we asked whether there is an interaction between SDC4 and VE-cadherin. We performed Proximity Ligation Assays (PLA) between SDC4 and VE-cadherin and could observe interactions between the two molecules under basal conditions (**Fig. 6A**). This interaction declined after 2 and 5 mins of VEGFA treatment and recovered after 10 minutes (**Fig 6B**). Quantitation of junctional SDC4/VE-Cadherin PLA spots saw a time dependent reduction in these interactions in response to VEGFA (**Fig. 6C**).

**Figure 6:**
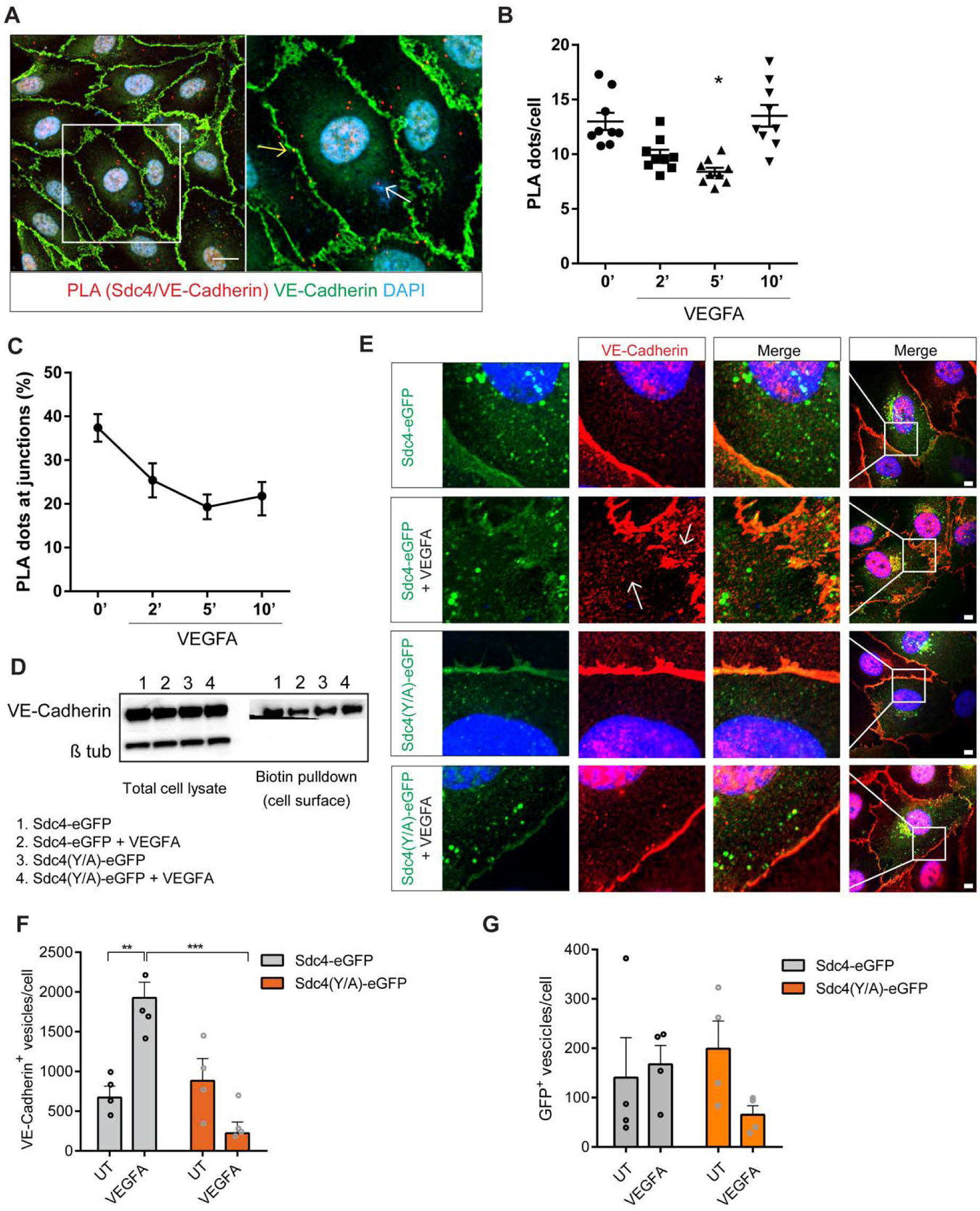
SDC4 associates with VE-Cadherin and its phosphorylation is required for VE-Cadherin internalization. **(A)** Confocal micrographs of HUVECs showing Proximity ligation on the cell surface (red dots) between HA tagged SDC4 and VE-Cadherin (nuclei, blue; VE-Cad, green). An example of a junctional dots (yellow arrow) and a non-junctional dots (white arrow) is shown on the right hand micrograph. Scale bar, 20 µm. **(B)** Quantification of SDC4HA/VE-Cadherin PLA puncta on the cell surface over time after VEGFA stimulation (10ng/ml). **(C)** Quantification of SDC4HA/VE-Cadherin PLA puncta at EC junctions over time after VEGFA stimulation (10ng/ml). **(D)** Cell surface VE-Cadherin is reduced after VEGFA stimulation in Sdc4-eGFP transfected cells but not in HUVECs transfected with Sdc4(Y/A)-eGFP. Cell surface biotinylation was performed on transfected HUVECs and biotin labelled proteins were ‘pulled-down’ using streptavidin beads. Lysates were analyzed by western blotting for VE-Cadherin. **(E)** Phosphorylated SDC4 is required for VE-Cadherin internalization. VE-Cadherin vesicles are evident in cells (white arrows) after VEGA stimulation in Sdc4-eGFP cells but not in Sdc4(Y/A)-eGFP treated cells, where VE-Cadherin remains at EC junctions. **(F)** Quantification of VE-Cadherin positive vesicles before and after VEGFA stimulation in HUVECs transfected with either Sdc4-eGFP or Sdc4(Y/A)-eGFP. (G) Quantification of eGFP+ vesicles (n=20 cells/condition).

In order to further investigate whether an interaction between SDC4 and VE-Cadherin was important for VEGFA driven VE-cadherin internalization, we first looked at the levels of cell surface biotinylated VE-Cadherin after VEGFA stimulation in HUVECs transfected with either Sdc4-eGFP or Sdc4(YA)-eGFP. In cells transfected with Sdc4-eGFP, a marked decrease in cell surface VE-Cadherin was observed after VEGFA treatment and this was not the case in cells transfected with Sdc4(Y/A)-eGFP (**Fig. 6D**). We then looked at the accumulation of VE-Cadherin containing vesicles in transfected HUVECs and noted a significant increase in intracellular VE-Cadherin^+^ vesicles in Sdc4-eGFP HUVECs when treated with VEGFA. However, in cells transfected with Sdc4(Y/A)-eGFP, vesicular trafficking of VE-Cadherin was greatly reduced (**Fig. 6E-F**).

### Soluble SDC4 reduces VE-Cadherin internalization and neovascularization in laser CNV

Lastly, we set out to explore whether a soluble form of SDC4 (referred to as solS4, corresponding to the complete ectodomain of SDC4) could interfere with its pro-angiogenic role, in particular by disrupting the SDC4/VE-cadherin interaction. Treatment of HUVECs with solS4 led to a reduction in PLA spots between endogenous SDC4 and VE-Cadherin (**Fig. 7A**). By using our antibody-feeding approach, we found that the addition of solS4 to a monolayer of WT ECs prevented VEGFA-driven VE-Cadherin internalization (**Fig.7, B and C**). In addition, solS4 treatment was also able to abolish VEGFA-driven EC migration; notably, and in contrast, ECs pre-treated with soluble SDC2 (solS2), the syndecan most structurally similar to SDC4, maintained responsiveness to VEGFA (**Fig. 7D**). Of note, the signaling kinases immediately downstream to VEGFR2 activation (i.e. Src, Erk and Akt) are not impacted by solS4 treatment, as evidenced by the increase of their phosphorylated active variants which is comparable between ECs treated with VEGFA and ECs treated with VEGFA and solS4 combined (**Fig. 7E and Fig. S10**). This evidence suggests that solS4 interferes directly with VE-Cadherin internalization step, bypassing the early VEGFA-driven signaling events. We also observed reduced sprouting when *ex vivo* aortic explants where treated with solS4 (**Fig. 7, F and G**). This led us to assess whether solS4 could be therapeutically beneficial as an anti-angiogenic compound, by testing its efficacy in the laser-induced CNV model in comparison to Aflibercept (Eylea®, Regeneron), one of the current standard therapies for neovascular AMD patients. We found that a single injection of solS4 at day 0 post-laser injury reduced the angiogenic response at day 7 by almost 50% compared to vehicle (PBS) control, achieving similar anti-angiogenic activity to that of Aflibercept (**Fig. 7, H and I**). These results indicate that delivery of solS4 decreases VE-Cadherin internalization, EC migration and reduces pathological angiogenesis in a murine model of neovascular AMD.

**Figure 7:**
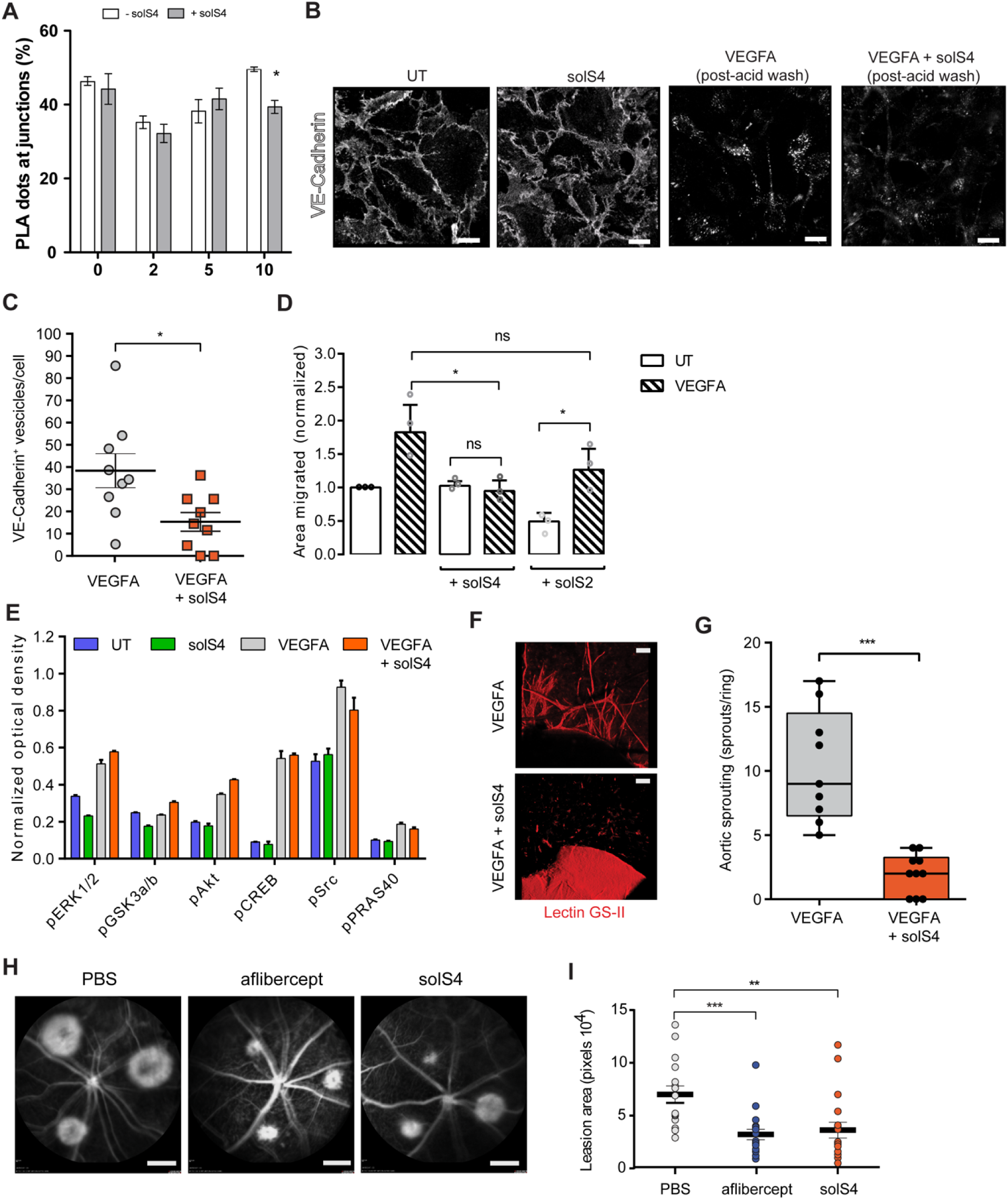
Soluble SDC4 reduces VE-cadherin internalization and neovascularization in laser CNV. **(A)** SolS4 disrupts PLA between SDC4 and VE-Cadherin at EC junctions. **(B)** Confocal images of VE-Cadherin antibody ‘washout’ experiments. WT MLECs were washed with PBS and incubated with anti-VE-Cadherin antibody at 4 °C for 1 hour. Unbound antibody was washed away and cells stimulated with VEGFA (30 ng/ml) for 10 min with or without SolS4 (3.5 nM) at 37 °C to promote VE-Cadherin internalization. Cells were then subjected to an acid wash to remove cell surface antibody. VE-Cadherin (white). Scale bar, 20 μm. **(C)** Images of MLECs treated with VEGFA and acid-washed were analyzed on Imaris software and VE-cadherin+ vesicles were counted and divided by the number of nuclei in the field of view. Images are representative of one experiment were 9 images per condition were analyzed. n=3 Scale bar, 20 μm. **(D)** HUVECs were scratched and incubated in serum-free media with or without VEGFA (20 ng/ml) and with or without SolS4 or solS2 (3.5 nM) for 16 hours. Migrated area was calculated by subtracting the final scratch area to the initial scratch area using ImageJ. n=3. **(E)** Confluent HUVECs were serum-starved for 2 hours and treated with soluble SDC4 (solS4, 3.5 nM), VEGFA (20 ng/ml) or both for 10 min. Lysates were then analyzed using a human phospho-kinase membrane-based sandwich immunoassay. Phosphorylated kinases were quantified using ImageJ, signals normalized using the reference spots. Phosphorylation sites were as follows, pERK1/2 (T202/Y204, T185/Y187), pGSK-3a,b (S21/S9), pAkt1/2/3 (S473), pCREB (S133), pSrc (Y419), pPRAS40 (T246). **(F and G)** Rings (1 mm in width) of aorta were dissected from adult WT rats, embedded in Collagen I and cultured as explants in VEGFA-containing medium with or without solS4 (3.5 nM) for 7 days and manually quantified. (n=4, 5-15 rings/condition). Scale bar, 10 μm. **(H)** Fundus fluorescein angiograms of WT animals at day 7 post laser induced CNV. Intravitreal injection of 1 µl of either PBS, Aflibercept (10 µg) or solS4 (100 ng) were performed at day 0 post-laser. Scale bar, 2.4 mm. **(I)** The CNV lesion areas were quantified using ImageJ, each dot represents the average of 3 lesions per eye. n=7-9 animals per condition. Data are means and error bars indicate SEM in B, C and G and min and max values in E. *P < 0.05; **P < 0.01; ***P<0.001.

## Discussion

As with other syndecan family members, *Sdc4-/-* animals show no gross abnormalities and develop normally, and, it is only after a challenge that phenotypes emerge (Corti et al, 2019; Echtermeyer et al, 2001; Ishiguro et al, 2000b; Johns et al, 2016). Consistent with this, we show that in disease models where pathological angiogenesis is a feature, *Sdc4-/-* animals are protected. Both tumor and ocular pathological angiogenesis are impaired in *Sdc4-/-* mice resulting in reduced tumor growth and reduced ocular neovascularization in the models used. We also show that SDC4 is the only family member whose expression is upregulated in a pathological scenario (murine OIR) and is present on newly formed, immature vessels in the retinas of diabetic retinopathy patients. Despite the defects in adult pathological angiogenesis, we observed no major defects in vascular development, other than a subtle phenotype during early retinal vascularisation. This is supported by our expression profiling data that showed SDC4 expression remains unaltered during developmental angiogenesis. These observations are consistent with other studies which show that developmental angiogenesis is not affected in the absence of SDC4, and it is in fact, the closely related SDC2 which is of importance for vascular development (Corti et al, 2019). Moreover, the existence of hypoxia-(Fujita et al, 2014) and inflammation-related regulatory elements (i.e. NF-κB (Okuyama et al, 2013)) within the *Sdc4* promoter region supports the concept that the expression and activity of this proteoglycan is driven by responses to hypoxia and inflammation, concomitant with pathological scenarios and important pro-angiogenic stimuli.

Reduced tumor volumes have been reported before, specifically in the Lewis Lung Carcinoma model and this correlated with elevated levels of NK cells in *Sdc4-/-* animals, the assumption being that these are responsible for the phenotype observed (El Ghazal et al, 2016). In both of the tumor models used in this study (papilloma and melanoma) analysis of immune infiltrates revealed negligible differences between WT and *Sdc4-/-* animals, suggesting that the NK cells phenotype might be Lewis Lung Carcinoma model specific. Additionally, in both models, vessel defects within the tumors were observed in *Sdc4-/-* animals consistent with defective tumor angiogenesis. VEGFA-stimulated angiogenic sprouting from *ex vivo* explants (choroid and aorta) was also reduced in the absence of SDC4, a process which has no leukocyte involvement. This, and the fact that angiogenesis in both tumor models is known to be VEGFA-driven supports the idea that the reduced tumor development seen in *Sdc4-/-* is due to impaired angiogenesis.

SDC4 has been shown be expressed in a wide variety of cell lines from many different tissues and is thought to be ubiquitously expressed albeit at low levels on all cell types (Kim et al, 1994). We revealed that SDC4 expression is upregulated during pathological angiogenesis in the eye and is selectively expressed on the ECs from immature blood vessels in diabetic retinopathy patients. VE-Cadherin is principally expressed on ECs and our findings relate specifically to the role of SDC4 in VEGFA-stimulated junctional disassembly which is a critical step in the formation of new vessels. Although likely, based on the above, we cannot exclude the possibility that the absence of SDC4 in other cell types may also be a contributory factor to the phenotypes observed. However, our observations using primary ECs from *Sdc4-/-* lungs that VE-Cadherin trafficking is attenuated would suggest that endothelial SDC4 is of major importance for normal pathological angiogenic responses.

Our initial hypothesis to explain the role of SDC4 in pathological angiogenesis was that SDC4 is involved in the interaction between VEGFA and VEGFR2 through interactions with its HS chains. Recent studies suggest that this is the case for VEGFC interactions with VEGFR3 during pathological lymphangiogenesis (Johns et al, 2016). However, analysis of *Sdc4-/-* ECs after stimulation with VEGFA revealed no reduction in either phosphorylation of VEGFR2 or major signaling kinases downstream of this receptor (Erk1/2), suggesting that in the absence of SDC4, VEGFA/VEGR2 signaling is unperturbed. These findings led us to speculate that SDC4 may have a role in angiogenesis downstream of the VEGFA/VEGFR2 axis. We demonstrated that SDC4 localizes to EC cell-cell contacts and co-localizes with the junctional protein VE-Cadherin. EC adherens junctions are a complex assemblage of both junctional and cytoskeletal proteins and many of the molecules involved also constitute focal adhesion complexes, in which SDC4 is a known component (Woods & Couchman, 1994). The disassembly of these structures is a critical early step in angiogenesis and we showed that ECs null for SDC4 exhibited defective VE-Cadherin trafficking away from junctions in response to VEGFA stimulation. Our data revealed that under basal conditions SDC4 is in complex with VE-Cadherin, and this complex is disrupted rapidly after VEGFA stimulation. The presence of SDC4 in this complex is clearly an important precursor for efficient VE-Cadherin trafficking since when SDC4 is absent (eg. in *Sdc4-/-* animals or ECs) this process is inhibited substantially. Likewise, when we expressed a non-phosphorylatable form of SDC4, in which the tyrosine residue (Y^180^) within the cytoplasmic domain is substituted with an alanine, VE-Cadherin internalization in response to VEGFA was significantly compromised. Interaction studies using FRAP suggest that this form of SDC4 is not in a molecular complex and this leads us to propose that under basal conditions it is the phosphorylated form of SDC4 which is interacting with VE-Cadherin and it is this that is required for efficient internalization. This raises the possibility that upon VEGFA stimulation SDC4 is dephosphorylated by a phosphatase leading to the disassembly of VE-Cadherin/SDC4 complexes and subsequent internalization of VE-Cadherin. This hypothesis is not unreasonable, since syndecan interactions with protein tyrosine phosphatases have been documented. For example, in joint synoviocytes SDC4 has been shown to modulate PTPRσ activity (Doody et al, 2015) and SDC2 has also been shown to interact with PTPRJ (CD148) (De Rossi et al, 2014; Tsoyi et al, 2018; Whiteford et al, 2011). Interestingly, the closest homologue to CD148 is VE-PTP and it is tempting to speculate that since SDC4 is the closest relative to SDC2 that this phosphatase may have a role in the interactions described. Alternatively, recent work demonstrated a role for SDC4 in calcium signaling in fibroblasts (Gopal et al, 2015) and this would also be consistent with the defects in VE-Cadherin redistribution we detect in *Sdc4-/-* ECs following treatment with VEGFA, given that the function of VE-Cadherin is calcium-dependent. Moreover, the expression of SDC4 has been recently shown to be required for the expression of junctional cadherin-11 in fibroblasts (Gopal et al, 2017).

Other studies have shown phosphorylation of the SDC4 cytoplasmic domain is important for SDC4 functionality. Phosphorylation of Y^180^ by c-Src has been shown to be a control point in integrin trafficking (Morgan et al, 2013) and c-Src is known to be involved in both VEGFR2 and VE-Cadherin signaling (Wallez et al, 2007). This suggests that c-Src is likely to be an essential component in the pathway described and may be responsible for the phosphorylation of SDC4 under basal conditions which allows it to complex with VE-Cadherin. Extensive structural studies have been performed on the SDC4 cytoplasmic domain and this reveals it to have a ‘twisted clamp’ structure (Koo et al, 2006; Lee et al, 1998). These studies also show that phosphorylation of S^179^ (adjacent to Y^180^) significantly alters the structure of the cytoplasmic domain and this was shown to have a profound impact on epithelial cell migration.

Our studies indicate that SDC4 is essential for efficient VE-Cadherin internalization in response to VEGFA and we show that the delivery of soluble SDC4 inhibits both VEGF-driven angiogenesis and hyperpermeability *in vivo*. From a therapeutic perspective, targeting of SDC4 with a view to inhibiting vascular permeability induced by VEGFA could offer avenues for the development of novel, selective therapeutics. Vascular leakage is an important and exacerbating factor in the early stages of diabetic retinopathy, not to mention the subsequent growth of new vessels associated with the later stages of this disease. This is also true of other neovascular eye diseases such as Wet AMD. The potential for blocking angiogenesis during tumor growth has been explored extensively with mixed results. Furthermore, the mechanism of resistance to the currently available anti-VEGF therapies both in tumors and retinopathy is actually related to the eradication of the neovessels, which, in turn, worsens the underlying ischemia and drives the formation of new, leaky blood vessels by alternative molecular mechanism. Targeting SDC4 would have the potential to block selectively pathological angiogenesis, but also could be used to stabilize tumor vessels to improve delivery of chemotherapy. As VEGFA signaling was not disrupted by the SDC4 inhibition, it could provide the survival signal for the ECs, but simultaneously reduce pathological permeability. Angiogenesis also occurs in the inflamed synovium of RA patients specifically in the formation of pannus and this also presents another potential target. Furthermore, SDC4 expression is up-regulated in RA and it mediates growth factor-/cytokine--induced inflammation in the disease (Cai et al, 2019). Anti-VEGF therapy is widely used in the successful treatment of eye diseases and some cancers (Gragoudas et al, 2004). Systemic application of these drugs can lead to side effects, not to mention the high level of patient non-response to these treatments. We show that SDC4 is downstream of VEGFA/VEGFR2 signaling and its expression is upregulated during pathological angiogenesis. It is tempting to speculate that targeting SDC4 to block angiogenesis/vascular permeability might actually have a less extensive side effect profile than anti-VEGFA therapies especially if administered systemically.

In summary, our findings identify SDC4 as an essential regulatory component in VEGFA-induced VE-cadherin trafficking from EC junctions during pathological processes and, therefore, provide significant further insight into the molecular events controlling neovascularization and hyperpermeability responses in diseases. The formation of abnormal blood vessels is a feature of cancer, neovascular eye diseases and chronic inflammatory diseases, which implies that SDC4 blocking strategies may have the potential to be applied in these contexts to either improve or offer a more selective alternative to existing therapies.

## Methods

### Study design

The objective of animal studies described here was to evaluate the role of SDC4 in developmental angiogenesis and in pathological neovascularization in the eye and to assess the feasibility of targeting SDC4 for treatment of neovascular eye diseases. Sample size was based on power calculations. In the laser induced CNV study, 7 represented the optimum number of animals needed to attain statistical significance of p<0.05 with a 90% probability given our pilot study values of 7 ±2 for group 1 and 4 ±1.5 for group 2. N was increased to 9 per group given the incidence of post-laser hemorrhage. The end-point of the experiment was day 7. Lesions that showed hemorrhage or two lesions fused together were excluded from analysis. For OIR experiments, 17 represented the minimum number of animals needed to attain statistical significance of p<0.05 with an 80% probability given our pilot study values of 4 ± 2 for group 1 and 2.5 ± 1 for group 2. N was increased to 21 per group. Three separate OIR experiments were performed and the end-point of the experiment was day 17. In developmental retinal angiogenesis study, 8 represented the optimum number of animals needed to attain statistical significance of p<0.05 with a 90% probability given our pilot study values of 30±4 for group 1 and 25±2 for group 2. N was increased to 9 per group. End-point of the experiment was day 6. Investigators were blinded during all phases of the experiments and genotypes or treatments groups only revealed after analysis.

### List of reagents

The following dyes and reagents were used in this work; lectin GS-II Alexa594 (Thermo Fisher Cat#L21416), DAPI (Sigma-Aldrich, Cat#D9542), Draq5 (Biostatus limited Cat#DR05500), recombinant mouse VEGFA 164 (R&D Systems Cat#493-MV-025m), Bradykinin (Sigma Cat#B3259), recombinant syndecan-4 (R&D Systems Cat# 6267-SD-050) and recombinant syndecan-2 (R&D Systems Cat#6585-SD-050), proteome profiler human phospho-kinase array kit (R&D Systems Cat#ARY003B). The antibodies used in these studies were as follows; IHC: anti-VEGFR2 (clone A-3 Santa Cruz cat#sc-6251)(Fig.1C and D), anti-VEGFR2 (cell signaling clone D5B1 Cat#9698 and Cat#2479) (Fig.5A), anti-Syndecan-4 (abcam cat#ab24511)(Fig.1C and D), anti-syndecan-4 (eBiovision cat#3644-100 lot 4AIIL36440)(Fig.5A), anti-CD31 (BD clone MEC13.3 cat#550274)(Fig.1C and D), anti-VE-Cadherin (abcam cat#ab33168)(Fig.4E), anti-VE-Cadherin (clone BV13 eBioscience Cat#14-1441-82)(Fig.4C, Fig.6A,Fig.S4A), anti-VE-Cadherin (Santa Cruz clone F-8 cat#sc9989 lot E1018)(Fig.5A and B), anti-α-SMA (1A4) (Sigma Cat#A2547)(Fig.3A and D, Fig.S3C and Fig.S4A). Western blotting: anti-VE-Cadherin (Santa Cruz clone F-8 cat#sc9989 lot E1018), anti-VEGFR2 (Cell Signaling clone D5B1 Cat#9698 and Cat#2479), anti-VEGFR2pY1175 (Cell Signaling clone D5B11 Cat#3770), anti-β tubulin (Sigma clone TUB2.1 lot 043M4785 Cat#T4026), anti-GFP (chromotek 3H9 lot60706001ab). Flow Cytometry: anti-VEGFR2-FITC (BD clone AVAS 12a1 lot 01750 Cat#560680), anti-IgG2a,k-FITC (BD Pharmingen Cat#553929), anti-Syndecan-4-PE (clone KY/8.2 lot 6204630 BD Pharmingen Cat#550352), anti-IgG2a,k–PE (BD Pharmingen Cat#553930), anti-VEGFR2 (AF644 89106 R&D System).

### Mice

*Sdc4-/-* mice were obtained from the Centre for Animal Resources and Development (CARD, Kunamoto University, Japan) with the kind permission of Professor Tetsuhito Kojima (Nagoya University, Japan). All adult mice were used at an age between 5-8 weeks at an average weight of 25 g. As appropriate, age-matched control mice were littermates or C57BL/6 wild-type mice purchased from Charles River. Animals were housed and treated in Accordance with UK Home Office Regulations and all experiments were approved by the UK Home Office according to the Animals Scientific Procedures Act 1986 (ASPA) and by the National Animal Ethics Committee of Finland.

### Skin tumor induction

*Sdc4-/-* and C57BL/6 WT mice were treated with DMBA and TPA to induce skin tumors according to the established protocol (Filler et al, 2007; May et al, 2015). In brief, the backs of 8 week old mice were shaved and 24 h later 50 µg DMBA (7,12-Dimethylbenz[a]anthracene) (Sigma, Dorset, UK) in 200 µl acetone was applied topically on the shaved area of the dorsal skin. After a week, the back skin of the mice was treated twice a week with 5 µg TPA (12-O-tetradecanoylphorbol-13-acetate) (Sigma) in 200 µl acetone for 21 weeks. Tumors (1 mm in diameter or larger) were counted twice a week. The fur excluding tumors was carefully shaved every two weeks.

### B16F1 melanoma tumor model

B16F1 mouse melanoma cells were obtained from HPA Laboratories and 5×10^7^ cells/ml in 100 μl of PBS were injected sub-cutaneously into the left flank of 6 week old WT and *Sdc4-/-* mice (n=12 per group). Mice were left for 7 or 14 days prior to sacrifice by cervical dislocation. A longitudinal incision was made from the chest to the genital area, skin was peeled and pinned down and tumors were revealed. Photographs were taken with a digital camera and tumors excised using scalpel and tweezers. Samples were then weighed on a precision balance and diameter measured using a ruler. Samples were then snap frozen in liquid nitrogen and stored at -80 °C. For analysis of melanoma immune infiltrates spleens and tumors were weighed, crushed through a 40 μm strainer, washed with 10 ml HBSS and incubated with ACK lysis buffer for 7 min at room temperature, washed again in HBSS and re-suspended in FACS buffer (PBS + 1 % NGS). Cells were counted, diluted to 10^6^ cell/ml in FACS buffer and 100 μl of suspension was incubated with 0.5 ug FcyIII/II R blocking antibody for 20 min at 4 °C. Subsequently, cells were stained with primary antibody solutions and incubated in the dark for 30 min at 4 °C. After 3 washes, samples were analysed using a BDLSR Fortessa cell analyser. Subsets of immune cells were identified using the following markers NK cells (NK1.1^+^, CD3^-^), Neutrophils (Gr1^high^, B220^-^CD3^-^), monocytes (Gr1^med/-^, B220^-^ CD3^-^), B cells (B220^+^, CD3^+^), T cells (CD3^+^, B220^-^), macrophages (F4/80^+^, B220^-^CD3^-^) and expressed as Subset % = No. events positive for subset marker/ No. events in leukocyte gate (for spleen samples) or Subset % = No. events positive for subset marker/ No. events in live cell gate (for tumor samples).

### Oxygen-induced retinopathy (OIR) mouse model

Neonatal mice at P7 were exposed to 75% oxygen for 5 days with their nursing mothers (Smith et al, 1994). At P12, they were returned to normoxia. Animals were euthanized at P12 to determine the area of vaso-obliteration or at P17 to determine the rate of retinal revascularization and pre-retinal neovascularization. As postnatal weight gain has been shown to affect outcome in the OIR model (Stahl et al, 2010), only weight matched (±1 g) pups were used in each experiment. Analysis of retinal vasculature as done as previously described (Vahatupa et al, 2016). Briefly, eyes were enucleated, fixed with 4% PFA for one hour and retinas dissected. Flat mount retinas were blocked in 10% normal goat serum and 10% fetal bovine serum for 2 hours, incubated overnight with isolectin GS-IB_4_ (1:100, Thermo Fisher). Briefly, retinas were imaged using confocal microscopy (Carl Zeiss LSM 700) with 10× objective. Pre-retinal neovascular tufts were readily distinguished from the superficial vascular plexus by focusing just above the inner limiting membrane. Areas of vascular obliteration and pathological neovascularization (neovascular tufts) were quantified using Adobe Photoshop CS3.

### Developmental angiogenesis mouse model

Neonatal WT and *Sdc4-/-* mice were euthanized at P6 and retinal flat mounts were prepared for lectin GS-II staining. Briefly, retinas were washed extensively in PBS, blocked in 3% BSA followed by overnight staining with lectin GS-II-Alexa594 and an anti-αSMA-alexa488 antibody (1:150, Sigma, in-house conjugated) in 0.5% triton in PBS. Retinas were imaged using confocal microscopy (Carl Zeiss LSM 700) with 10× objective. The rate of developmental angiogenesis at P6 was determined by measuring the diameter of vasculature via the optic nerve to the tips of the blood vessels. Two measurements per retina were taken and an average was calculated.

### Laser-induced choroidal neovascularization (CNV) mouse model

WT and *Sdc4-/-* 6 week old mice were anesthetized by intraperitoneal injection of 0.15 ml of a mix of Domitor and Ketamine and pupils dilated with topical administration of 1 % Tropicamide and 2.5 % Phenylephrine. Three burns per eye were be made by laser photocoagulation (680 nm; 100 μm spot diameter; 100 ms duration; 210 mW). Only burns that produced a bubble, indicating the rupture of the Bruch’s membrane, were counted in the study. At day 7 after the laser injury, fundus fluorescein angiography (FFA) was performed to measure the area of CNV lesion. The area of lesion was then quantified using Imaris Software (Bitplane) and expressed as number of pixels. Additionally, at the end of the experiment (day 7) mice were culled, choroid-RPE tissue dissected, flat-mounted and stained with lectin GS-II. Confocal images were then acquired using a PASCAL laser-scanning confocal microscope (Carl Zeiss) followed by volumetric analysis with Imaris software (Bitplane).

### Human samples

Pre-retinal neovascular membranes were obtained from type I diabetic patients who were undergoing pars plana vitrectomy for the treatment of proliferative diabetic retinopathy. These patients had already developed tractional retinal detachment due to fibrosis of neovascular membranes. Thus, the samples represent the end stage disease, where substantial amount of fibrosis was associated with neovessels, but still contained regions with active pathologic angiogenesis. All patients were Caucasians, males and females, aged between 27–56 years, and mean duration of diabetes was 24 years (range, 21–28 years). The protocol for collecting human tissue samples was approved by the Institutional Review Boards of the Pirkanmaa Hospital District. The study was conducted in accordance with the Declaration of Helsinki. All patients gave written informed consent. During vitrectomy, the fibrovascular membranes were isolated, grasped with vitreous forceps, and pulled out through a sclerotomy. The sample was immediately fixed with 4 % formaldehyde for 3 h, transferred to 70 % ethanol, embedded in paraffin, and processed for immunohistochemistry. Adjacent 5 µm sections were stained for anti-CD31, anti-VEGFR2 and anti-SDC4 KY/8.2 followed by horseradish peroxidase (HRP) conjugated secondary antibodies. The slides were imaged via Olympus light microscope and Cell Sens Dimensions software. Quantification was done using IHC Profiler plugin in ImageJ 1.44p software (National Institutes of Health, Bethesda, MD). The same region of interest was selected from all stained sections (2-3 sections per sample) and the area (%) of total positive staining was quantified. To analyze the correlation between SDC4, VEGFR2 and CD31 positive area in each ROI, a multiple linear regression analysis was done in IBM SPSS Statistics.

### Evans blue permeability assay

6-8 week old mice were anesthetized by i.m. injection of 1 ml/kg ketamine (40 mg) and xylazine (2 mg) in saline solution. The back skin was shaved using an electric razor. Mice then received Evans Blue dye (0.5 % in PBS, 5 μl per g bodyweight) i.v. through the tail vein. Afterwards, 50 µl of PBS containing either 100 ng of VEGFA or 100 μg of Bradykinin or PBS alone were injected s.c. in the mouse dorsal skin. After 90 min animals were sacrificed by cervical dislocation. Dorsal skin was removed and injected sites were cut out as circular patches using a metal puncher (∼8 mm in diameter). Samples were then incubated in 250μl of formamide at 56 °C for 24 h to extract Evans Blue dye from the tissues. The amount of accumulated Evans Blue dye was quantified by spectroscopy at 620 nm using a Spectra MR spectrometer (Dynex technologies Ltd., West Sussex, UK). Results are presented as the optical density at 620 nm (OD620) per mg tissue and per mouse.

### Choroid and aortic sprouting assay

Aortic rings were prepared from 6-8 week old male mice. Briefly, aortas were dissected, washed in PBS and fat and side branches were removed prior to slicing into rings (∼1mm diameter) and these were incubated o/n in serum-free OptiMEM (containing Penicillin-Streptomycin) at 37 °C. Choroid explants were prepared from eyes, as follows; eyes were punctured with scissors, an incision was made 1 mm below the iris and the iris/cornea/lens/retina were removed. The choroid was cut into approximately 1 mm^2^ pieces (∼ 4 from central, 6 from periphery) and incubated o/n in OptiMEM, as above. Explants were embedded in 150 µl/well of a type I collagen matrix (1 mg/ml) in E4 media (Invitrogen) in a 48-well plate (Corning). Plates were incubated at 37°C for 30 min to allow the collagen to solidify, after which time wells were supplemented with 200 µl OptiMEM with 1 % FBS and VEGFA (30 ng/ml) or FGF (10 ng/ml) or both and incubated at 37°C, 10 % CO2. Media was replenished every third day and angiogenic sprouts from aortic rings or choroid explants were counted after 1 week and results were expressed as the number of sprouts per ring or explant.

### Matrigel plug assay

Matrigel (400 µl, BD Biosciences) was thawed on ice and mixed with 100 μl PBS containing growth factors 100 ng/ml VEGFA and 20 U/ml of Heparin (Sigma) and injected subcutaneously into the flank of anesthetized 6 week old mice roughly between the rib cage and the posterior leg. After 5 days, mice were sacrificed by cervical dislocation and plugs recovered after dissection. Plugs were imaged using a conventional digital camera, weighed on a precision balance and incubated overnight at 4 °C with 500 μl of dH2O. The amount of haemoglobin released from the plugs was measured using the Drabkin reagent kit (Sigma) as described by the manufacturer. Absorbance at 540 nm was read using a spectrophotometer and results quantified using a standard curve of known concentrations of haemoglobin and data was expressed as the concentration of haemoglobin per gram of plug (HG mg/ml/g).

### Primary mouse lung endothelial cell (MLEC) isolation and HUVEC culture

Cells were isolated and cultured as described (Reynolds et al, 2002). HUVECs were obtained from Promocell and were grown and maintained according to supplier’s instructions. RAW 246.7 macrophages were obtained from HPA culture laboratories (Public Health Laboratories, UK) and grown in DMEM supplemented with 10% FCS.

### Syndecan-4 eGFP and HA tagged lentiviral constructs

cDNA encoding the full length sequence of murine SDC4 (Sdc4-eGFP) or SDC4 mutated at position Y^180^ (Sdc4(Y/A)-eGFP) were synthesized to include the complete coding sequence of eGFP inserted between I^32^ and D^33^ of the extracellular domain. An HA tagged version of murine SDC$ was also synthesized with the coding sequence being inserted in between the same amino acids (GeneArt, Life Technologies). These cDNAs were ligated into the BamHI and XhoI sites of the lentiviral vector plentiSFFV-MCS using conventional procedures. Lentiviruses were packaged in HEK 293T cells under the control of the GagPol promoter using the VSVG envelope protein. HUVECs were transfected by overnight incubation with viral preparations.

### Migration assay

Cells were cultured in 6- or 12-well plates until confluent and serum starved for 3 hours. One longitudinal scratch was introduced to the monolayer with a sterile pipette tip, cells were then washed to remove debris and incubated in fresh medium. Regions of interests in the scratch area were recorded using Cell^M software (Olympus) and micrographs of ROIs taken every 30 min for 16 hours with an Olympus IX81 inverted microscope. The initial and final gap was measured using ImageJ software (NIH).

### VE-Cadherin internalization assay

Cells were serum starved for 3 hours prior to incubation for 1 hour at 4 °C in the presence of an anti-VE-Cadherin antibody (clone BV13, eBioscience) conjugated in-house to Alexa555. Unbound antibody was washed away using serum free medium. 30 ng/ml of VEGFA was added to cells for 5 min at 37 °C to allow internalization of VE-Cadherin and bound antibody. For analysis of internalized VE-Cadherin, cell surface bound antibody was removed using an acid wash (glycine 25 mM, 3% BSA in PBS, pH 2.7). Cells were washed 3X with ice cold PBS and fixed with 4 % PFA, 0.25 % glutaraldehyde prior to analysis by confocal microscopy.

### Biotinylation assay

Cells were serum starved for 3 hours prior to incubation with VEGFA (50 ng/ml) for 5 min. Cells were washed 3 times with PBS (Ca^+^, Mg^+^) on ice and incubated with 1:500 v/v biotin (sulfo-NHS-biotin, thermo fisher, 12.5 ng/µl) in PBS for 20 min also on ice andwashed twice in ice-cold PBS, blocked for 20 min in complete medium and lysed (TBS, 10 mM EDTA, 1% Triton, phosphatase inhibitor cocktail (Halt, thermos fisher)). Lysates were centrifuged at 14000 rpm for 5 min and supernatant collected (this will be total protein). A fraction of the lysate was then incubated with Neutravidine-agarose (thermos fisher) for 2 hours at 4 °C following 3 washes in lysis buffer by pulse centrifugation. Samples were then diluted in 3X laemmli buffer and heated at 65 °C for 15 min and analysed by western blotting.

### Proximity ligation assay

Proximity ligation assay or PLA is a technique that allows the identification in situ of spots in which 2 proteins are in close proximity (less than 40 µm). The PLA experiments were performed as per manufacturer’s instructions (Duolink, Sigma Aldrich). Cells were treated as indicated in the results section. Following treatments, medium was taken off, cells were washed in PBS at RT, fixed in 4 % PFA for 15 min at RT. After a 5 min wash in PBS, PFA was quenched by incubating cells with 0.1 M NH4CL for 10 min at RT.

### qPCR

RNA was purified using the Qiagen RNeasy Microkit as described by the manufacturers, cDNA was synthesized from 10 ng total RNA using the iScript cDNA synthesis kit (Biorad), quantitative real time PCR was performed on an ABI7900HT (Applied Biosystems) real time PCR machine using the iQ Sybr green supermix (Biorad). Sequences of primers used in this are as follows: Mouse syndecan-1 forward 5’-GGT GGA CGA AGG AGC CAC A-3’, mouse syndecan-1 reverse 5’-CTC CGG CAA TGA CAC CTC C-3’, mouse syndecan-2 forward 5’-GAG GCA GAA GAG ATG CTG G-3’, mouse syndecan-2 reverse 5’-CAT CAA TGA CGG CTG CTA G-3’, mouse syndecan-3 forward 5’-GCC CTG GCC TCC ACG AC-3’, mouse syndecan-3 reverse 5’-CAC GAT CAC GGC TAC GAG C-3’, mouse syndecan-4 forward 5’-CAC GAT CAG AGC TGC CAA G-3’, mouse syndecan-4 reverse 5’GGA TGA CAT GTC CAA CAA AGT-3’, mouse GAPDH forward 5’-TCG TGG ATC TGA CGT GCC GCC TG-3’, mouse GAPDH reverse 5’-CAC CAC CCT GTT GCT GTA GCC GTA T-3’.

### Statistical analysis

Data are presented as mean ± s.e.m. and sample sizes are reported in each figure legend. Each experiment on animals was performed on at least two independent litters of a given genotype. Data were plotted and analyzed for statistical significance using Prism 6 (Graphpad software Inc.). Parametric statistical tests (two-tailed t-test and one- or two-way ANOVA) were used to compare the averages of two or more groups, with post-hoc Tukey’s or Bonferroni’s multiple comparison tests. P-values less than 0.05 were considered statistically significant.

## Supplementary Materials

Supplementary materials and Methods

Fig. S1. B16F1 melanoma cells express SDC4.

Fig. S2. Flow cytometric analysis of immune cells in B16F1 melanomas are the same between WT and *Sdc4-/-* animals.

Fig. S3. Infiltrated Immune cells in the DBA/TMPA model are the same between WT and *Sdc4-/-* animals.

Fig. S4. Flow cytometric analysis of the immune cell profiles in spleen and skin draining lymph nodes (LN) form WT or *Sdc4-/-* animals after 21 weeks of DMBA/TPA treatment.

Fig. S5. Cell proliferation and skin thickness are the same between WT and Adc4-/- animals in the DMBA/TPA model.

Fig. S6. *Sdc4-/-* response to hyperoxia is comparable to WT.

Fig. S7. Developmental retinal angiogenesis in WT and *Sdc4-/-* mice.

Fig. S8. Analysis of adult vascular beds in WT and *Sdc4-/-* mice.

Fig. S9. VEGFR2 and Sdc4-eGFP expression in transfected HUVECs.

Fig S10. solS4 does not affect VEGFA/VEGFR2 signalling. References.

## Acknowledgments

**Funding**: JRW and GDR gratefully acknowledge funding from Arthritis Research UK (Grant No. 19207 and 21177), Fight for Sight (Grant No. 1558/59), Barts and The London Charity (Grant No. MGU0313), Queen Mary Innovations, William Harvey Research Foundation and The Macular Society. TAHJ, MV and HU-J gratefully acknowledge funding from the Academy of Finland, Päivikki and Sakari Sohlberg Foundation, Instrumentarium Research Foundation, Diabetes Wellness Foundation (DWF), Pirkanmaa Hospital District Research Foundation, Tampere Tuberculosis Foundation and the Finnish Cultural Foundation. JWB is a NIHR Research Professor.

## Author contributions

GDR, JRW, EC and TAHJ designed the experiments; GDR, JRW, EC, TAHJ, MV, HU-J interpreted the results; GDR and JRW wrote the manuscript; GDR performed choroid and aortic ring assay, matrigel plug assay, lentivirus production (JRW designed the cDNAs), cell migration assay, VE-Cadherin internalization assay, permeability assay, immunofluorescence staining, biotinylation experiments, vascular beds staining, SDS-Page and western blotting, flow cytometry analysis; EC, GDR, SEL performed the laser induced CNV and in conjunction with MV and HU-J performed the analysis of neonatal retinal development; MV, HU-J, TAHJ performed the OIR experiments and staining. MV and HU-J performed the staining and analysis of human PDR membranes. We thank Marianne Karlsberg and Terhi Tuomola for practical support.

## Data availability Statement

The datasets generated during and/or analysed during the current study are available from the corresponding author on reasonable request.

